# HIV-specific T-cell responses reflect substantive in vivo interactions with infected cells despite long-term therapy

**DOI:** 10.1101/2020.08.28.272625

**Authors:** Eva M. Stevenson, Adam R. Ward, Ronald Truong, Allison S. Thomas, Szu-Han Huang, Thomas R. Dilling, Sandra Terry, John K. Bui, Talia M. Mota, Ali Danesh, Guinevere Q. Lee, Andrea Gramatica, Pragya Khadka, Winiffer D. Conce Alberto, Rajesh T. Gandhi, Deborah K. McMahon, Christina M. Lalama, Ronald J. Bosch, Bernard Macatangay, Joshua C. Cyktor, Joseph J. Eron, John W. Mellors, R. Brad Jones, for the ACTG A5321 Team

## Abstract

Antiretroviral therapies (ART) durably suppress HIV replication to undetectable levels – however, infection persists in the form of long-lived reservoirs of infected cells with integrated proviruses, that re-seed systemic replication if ART is interrupted. A central tenet of our current understanding of this persistence is that infected cells are shielded from immune recognition and elimination through a lack of antigen expression from proviruses. Efforts to cure HIV infection have therefore focused on reactivating latent proviruses to enable immune-mediated clearance, but these have yet to succeed in driving reductions in viral reservoirs. Here, we revisited the question of whether HIV reservoirs are predominately immunologically silent from a new angle, by querying the dynamics of HIV-specific T-cell responses over long-term ART for evidence of ongoing recognition of HIV-infected cells. We show that T-cell responses to autologous reservoir viruses persist over years, and that the maintenance of HIV-Nef-specific responses was uniquely associated with residual frequencies of infected cells. These responses disproportionately exhibited a cytotoxic, effector functional profile, indicative of recent *in vivo* recognition of HIV-infected cells. These results indicate substantial visibility of the HIV reservoir to T-cells on stable ART, presenting both opportunities and challenges for the development of therapeutic approaches to curing HIV infection.

## Introduction

The needs for both a vaccine and a cure for HIV are underscored by the ongoing impact of this global pandemic, which continues to cause close to 800,000 deaths annually (1). Antiretroviral therapy (ART) is capable of durably suppressing HIV replication, and halting disease progression for those able to access and adhere to these regimens. Infection persists, however, in reservoirs of CD4^+^ T-cells, and potentially other cell types (2, 3) with integrated proviruses that re-seed systemic replication if ART is interrupted (2, 4–9). These proviruses often exist in a latent state, characterized by limited transcription and, presumably, a lack of antigen production. This gives rise to one of the central tenets in the study of HIV persistence, which postulates that the persistent reservoir (often called the ‘latent reservoir’) is not detected by the immune system in individuals on long-term ART. It follows that engaging the immune system to reduce HIV reservoirs depends upon latency reversal to re-expose the immune system to HIV antigen – the so-called “kick and kill” (or “shock and kill”) strategy (10).

While latency undoubtedly diminishes immune recognition of viral reservoirs, several lines of evidence cast doubt on whether this is absolute *in vivo*, which would implicate additional contributors to viral persistence (11). Most notably, unspliced, and sometimes multiply spliced, HIV transcripts are readily detectable in peripheral blood mononuclear cells (PBMCs) of individuals on durable ART (12, 13). These observations have recently led some to propose amendments to the “latent reservoir” model, by introducing the idea of a continuum ranging from “deep latency” (no RNA produced) through to an “active reservoir” (14, 15). A key unresolved question, however, is whether these transcripts result in HIV-protein production, and thus enable immune recognition. Multiple factors limit the degree to which this can be inferred from direct measures of *in vivo* viral expression, including sampling difficulties – given that expression may be anatomically or temporally restricted – and the lack of equivalency between readily measurable features (ex. viral RNA) with bonafide antigen presentation (16, 17). We therefore hypothesized that some level of antigen recognition by HIV-specific T-cells may occur *in vivo* in ART-suppressed individuals with undetectable viremia. We predicted that this would be reflected in relationships between the long-term dynamics of HIV-specific T-cell responses and measures of virologic persistence, including frequencies of infected cells.

Although the T-cell response to HIV infection has been generally well characterized, and is known to decay rapidly in the months following ARV initiation (18–20), there are a lack of well-powered studies that have addressed the long-term dynamics of these responses in association with virologic parameters. In a previous cross-sectional study, we observed a modest correlation between the magnitudes of T-cell responses to the HIV-Nef protein and residual frequencies of infected cells, providing some initial suggestion that these responses may be maintained by antigen recognition. However, a recent longitudinal study reported that, while responses were highly stable on durable ART, no correlations were observed between response magnitudes and reservoir size as measured by quantitative viral outgrowth assays across 18 individuals (21). The current study builds upon these earlier reports by uniquely assessing T-cell response dynamics over almost 3 years in association with multiple measures of viral persistence, in a cohort of 49 individuals on well-documented sustained ART. We first confirm that, in this cohort, T-cell responses to autologous reservoir viruses are well represented by a scalable IFN-γ enzyme-linked immunospot (ELISPOT) assay, and show that these responses persist over years. Strikingly, the persistence of T-cell responses to the HIV-Nef protein (slopes of change) over 144 weeks were strongly and uniquely associated with the frequencies of infected cells that persisted on ART (22, 23), and these responses disproportionately exhibited a cytotoxic effector functional profile, indicative of recent *in vivo* antigen recognition (24–28). These results conclusively reveal ongoing interactions between the immune system and the HIV reservoir over years of ART, with implications both for understanding HIV persistence, and designing interventions aimed at curing infection.

## Results

### CD8^+^ T-cell Responses to Autologous Infected Cells

We approached the characterization of CD8^+^ T-cell responses in our study with initial concerns over putative limitations in conventional approaches to quantifying T-cell responses to cells infected with autologous reservoir viruses. Namely, by utilizing synthetic peptides as antigens, conventional approaches may: i) detect responses from T-cells that are unable to recognize autologous reservoir viruses, as a result of sequence variation (13, 29); ii) not fully capture the entirety of viral epitopes, which may also be expressed from cryptic reading frames, or novel exon structures (23); and iii) skew representation of epitopes that are differentially affected by processing in infected cells (30).

With the aim of more comprehensively quantifying the total ability of CD8^+^ T-cells to recognize infected cells, we developed a ‘biosensor assay’ whereby *ex vivo* CD8^+^ T-cells were co-cultured with excess HIV-superinfected autologous CD4^+^ T-cells. For each individual (participant characteristics in **Table S1**), we prepared two sets of target cells infected with either: i) the molecular clone of HIV, JRCSF, or ii) a cocktail of autologous reservoir viruses generated by pooling the supernatants of quantitative viral outgrowth assays (**Fig. 1A & B**). Flow cytometric analysis detected CD8^+^ T-cells that responded to these infected cells by degranulating (CD107a) and/or producing IFN-γ, in most individuals (**Fig. 1C**). In comparing these biosensor assay responses (to autologous viruses) with the total IFN-γ ELISPOT responses (summed across all HIV gene products), we observed a strong correlation (Spearman r=0.840, p=0.005, **Fig. 1D**), in spite of the above-noted putative limitations with using synthetic peptides. Thus, these data from our ‘biosensor assay’ serve to not only directly demonstrate that CD8^+^ T-cell responses capable of recognizing cells infected with autologous reservoir viruses remain present in individuals on long-term ART, but also to show that such responses are reasonably well represented by ELISPOT results, when summed across all HIV gene products.

**Fig. 1.**
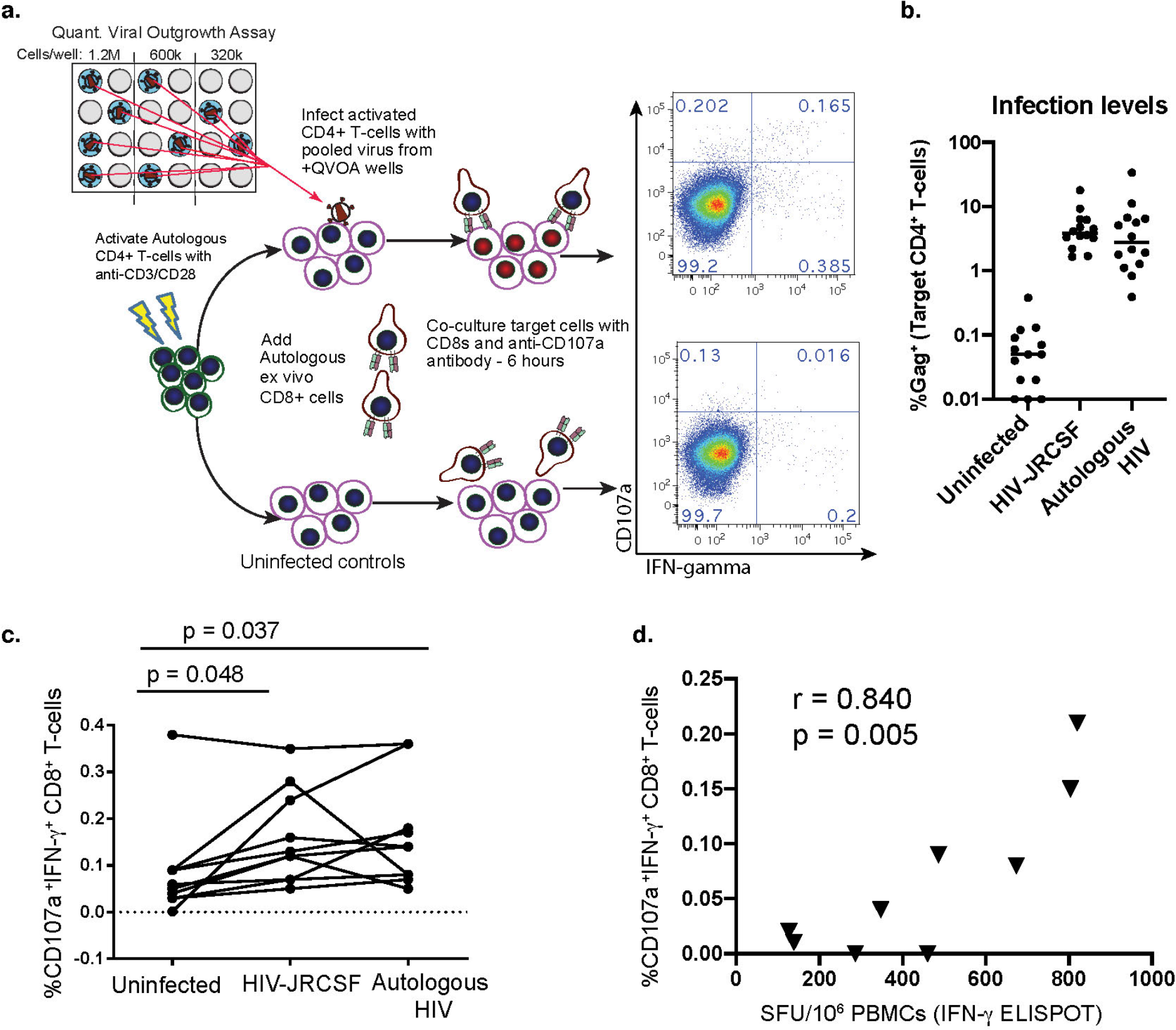
CD8^+^ T-cell responses to virus-infected cells can be detected *ex vivo*, correlating with ELISPOT responses. **A.** Schematic of ‘biosensor assay’. Top-right: Flow cytometry plot gated on CD8^+^ T-cells co-cultured with infected CD4^+^ T-cells. Bottom-right: Uninfected control. **B.** CD4^+^ target cells’ infection levels from all assays, measured by flow cytometry. **C.** Flow cytometry data depicting %CD107a^+^IFN-γ^+^ of viable CD8^+^ T-cells, each line representing a different participant (means from 3 replicates). P-values calculated by RM one-way ANOVA with Dunnett’s multiple comparison test. **D**. Spearman’s correlation between background (uninfected) subtracted responses to autologous HIV (as in **C**), and summed IFN-γ ELISPOT responses to HIV proteome.

### Magnitudes of T-cell Responses on Long-Term ART

With the above validation in place, we leveraged the scalability of the ELISPOT assay to comprehensively examine T-cell response dynamics in a larger cohort. These assays were performed using overlapping 15-mer peptides spanning: i) HIV-Gag, ii) HIV-Env, iii) HIV-Pol, iv) HIV-Nef/Tat/Rev, v) HIV-Tat, vi) HIV-Rev, vii) HIV-Nef, and viii) CMVpp65 (control), with samples from the ACTG A5321 HIV Reservoirs Cohort Study, which consists of participants who initiated ART during chronic HIV infection and had subsequent well-documented, sustained virologic suppression (undetectable by clinical assay prior to and throughout the study period) (31) (**Fig. 2 and Table 1**). We previously assessed HIV-specific T-cell responses in A5321 at study entry, a median of 7 (range 4-15) years after ART initiation (32). Here, we extended these results with batched analysis of samples from 24 and 168 weeks after study entry. IFN-γ-producing HIV-specific T-cell responses were readily detected against Gag, Pol, and Nef, with median values at 24 weeks: 103, 78.5, and 78.5 SFU/10^6^ PBMCs, respectively; and at 168 weeks: 87.0, 44.7, and 43.3 SFU/10^6^ PBMCs, respectively (**Fig. 3A&B and Table S2**).

**Fig. 2.**
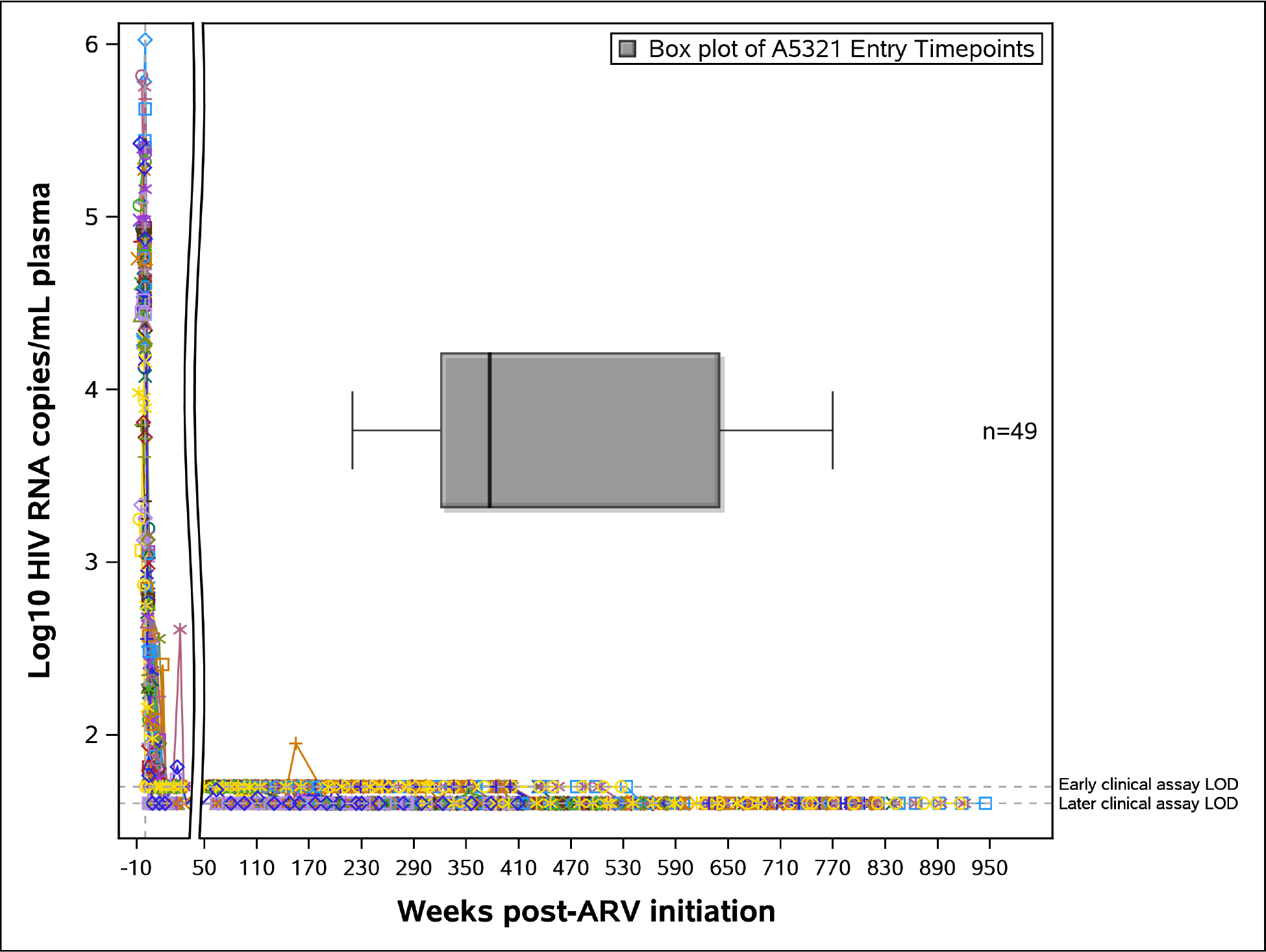
ACTG A5321 Cohort participants achieved viral suppression prior to study entry and maintained viral suppression throughout the study period. Log_10_ plasma HIV RNA (copies/mL) by clinical commercial assays for ACTG A5321 Cohort study participants included in this longitudinal sub-study, followed from pre-ART initiation (ART initiated in other ACTG trials) through to the A5321 study 168 week timepoint. Limit of detection (LOD) for early clinical assays was 50 copies/mL, and for later clinical assays 40 copies/mL. Colored lines represent individual participants (n=49), with symbols indicating each clinical viral load measurement. X-axis break shows time post-ART initiation when all participants achieved initial viral suppression. Box plot shows the distribution of participants’ A5321 study entry timepoints relative to weeks post-ART initiation (minimum, Q1, median, Q3, maximum).

**Table 1.**
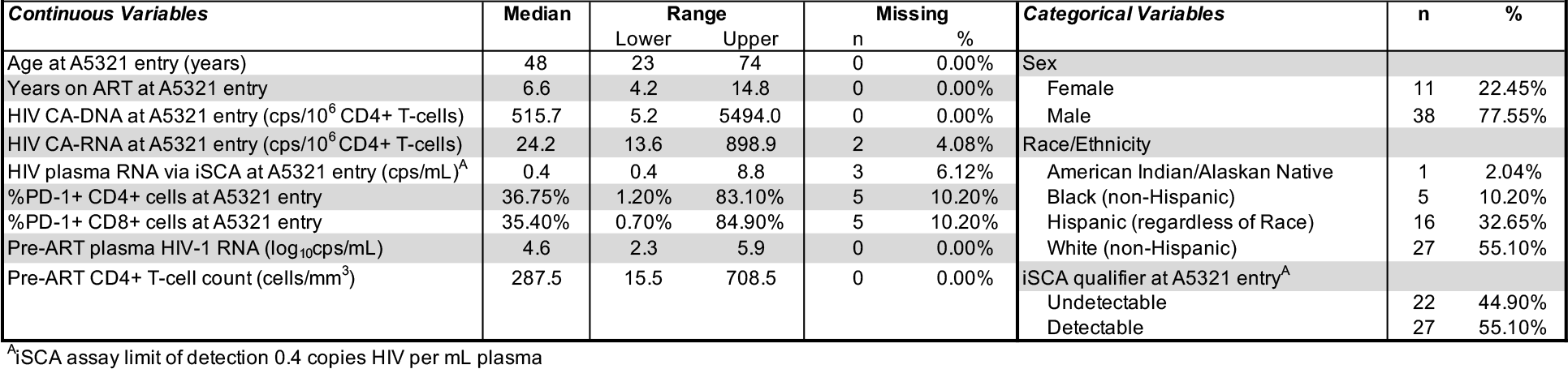
Demographic, virologic, and immunologic characteristics of longitudinal sub-study participants

**Fig. 3.**
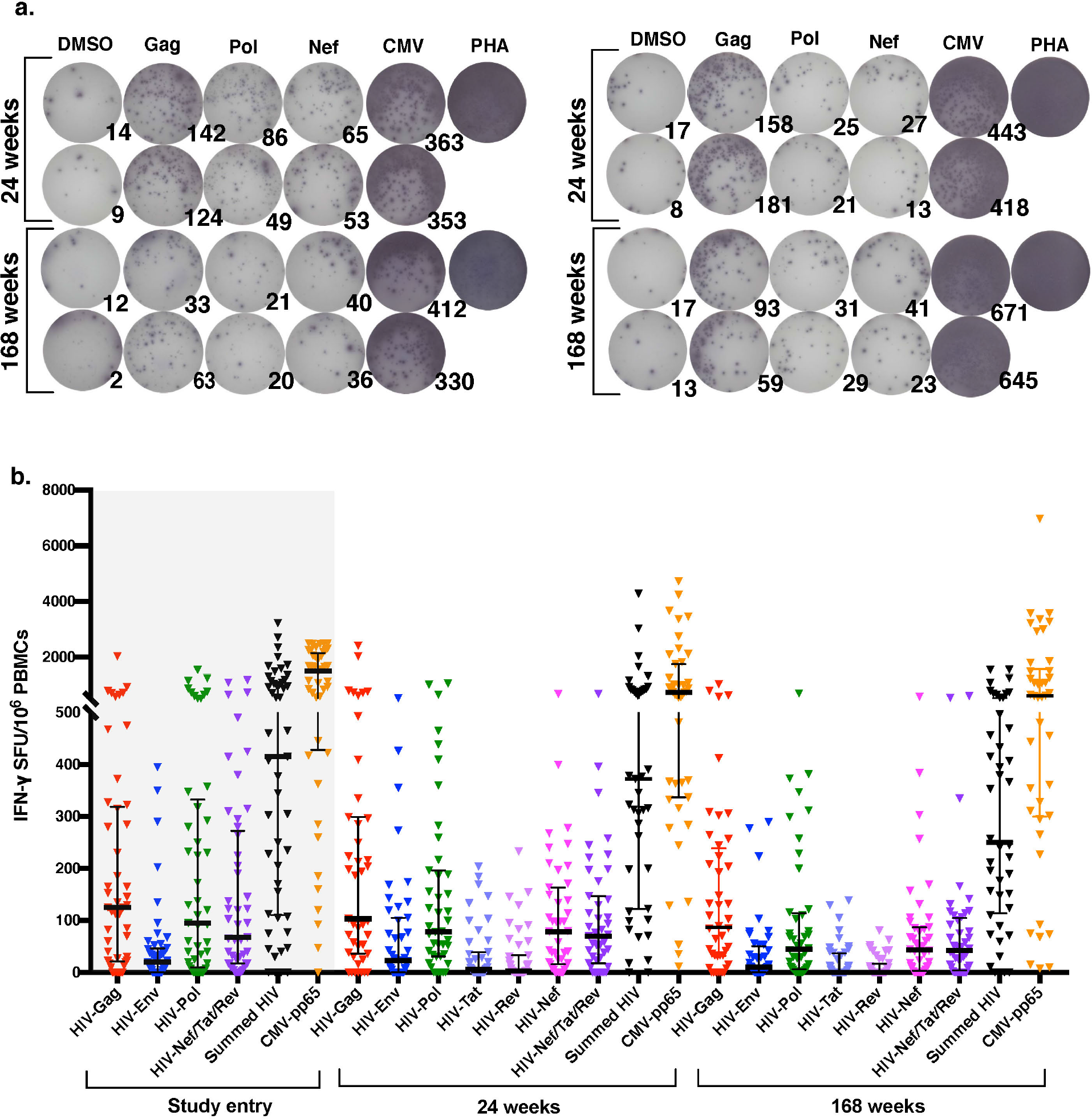
HIV-specific T-cell responses readily detectable *ex vivo* and persist on long-term ART, primarily directed against HIV-Gag, HIV-Pol, and HIV-Nef. **A.** Representative IFN-γ ELISPOT results for two participants for both timepoints, with 2×10^5^ PBMCs/well. **B.** Magnitudes of IFN-γ responses are shown for three on-ART timepoints. Study entry timepoint data is shaded in gray because it was not performed in batch with 24 and 168 weeks timepoints. Each data point represents the mean spot forming units (SFU)/10^6^ PBMCs following background subtraction of negative control wells (duplicates). Vertical lines and error bars represent median and interquartile range for each peptide pool.

Between this 24 to 168 week period, time-averaged responses against Gag were the highest, and significantly greater than responses to Env, Nef, Tat, and Rev (all p<0.05) (**Table S3**). Notably, T-cell responses directed against Tat and Rev were the lowest in magnitude, and negligible at both timepoints (**Fig. 3B and Tables S2 & S3**), Env-specific responses were also low, with median values of 22.9 and 10.3 SFU/10^6^ PBMCs at 24 and 168 weeks, respectively (**Fig. 3B and Table S2**). The long-term persistence of HIV-specific T-cell responses – primarily directed against Gag, Pol, and Nef – over years of ART thus provided initial support for these HIV-specific T-cells continuing to interact with persisting infected cells.

### Maintenance of Nef-Specific T-cells by the Reservoir

To further characterize the long-term dynamics of HIV-specific T-cell responses in A5321 cohort participants on durable ART, we categorized participants’ IFN-γ ELISPOT responses from the batched 24 to 168 weeks post-study entry data as either increasing, decreasing, or not changing (defined as ≤15% change), and observed considerable heterogeneity (**Fig. S1**). Notably, population-average responses to Nef, summed HIV, and CMV-pp65 did not decline significantly over this 144 week time period, whereas responses to Gag, Env, and Pol all showed significant declines over time (**Fig. 4A and Table S4**). However, all HIV-specific T-cell responses demonstrated remarkable persistence, with the responses which showed a significant decline only averaging between 0.35% to 0.62% loss per week in IFN-γ ELISPOT assays (**Table S4**).

**Fig. 4.**
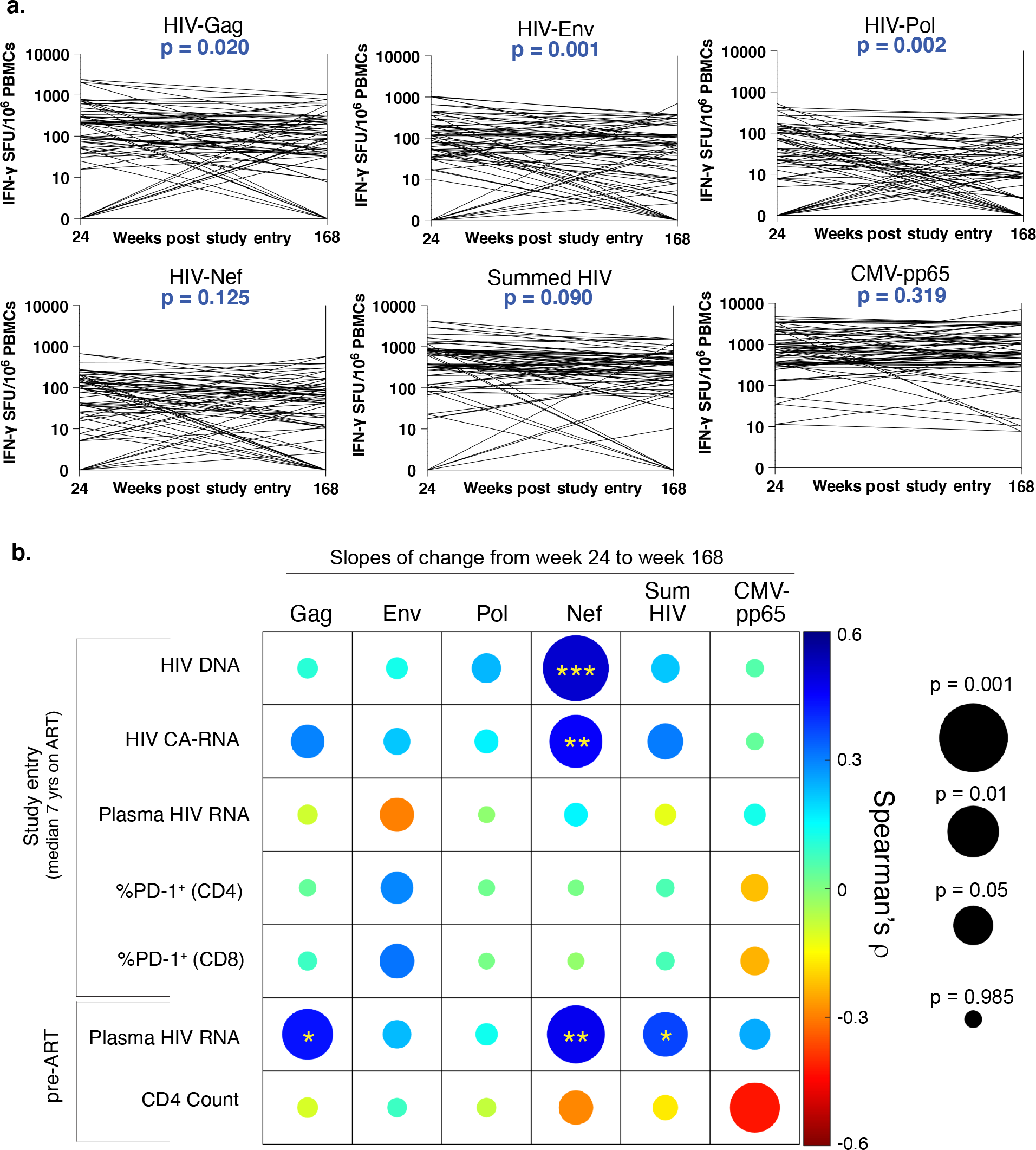
HIV-specific T-cell responses highly stable on long-term ART, with HIV-Nef-specific response dynamics uniquely associated with reservoir measures. **A.** Participant-specific slopes of change in T-cell responses from weeks 24 to 168 post-study entry. P-values represent the significance level for the covariate time (in weeks) in linear mixed-effects models from **Table S4. B.** Correlogram depicting Spearman correlations between slopes of change in raw magnitudes of T-cell responses (from panel **A**) with virologic and immunologic parameters. Color scale bar represents magnitude of correlation coefficient. Circle size represents unadjusted p-values. Asterisks represent adjusted p-values controlling for pre-ART plasma HIV RNA and CD4^+^ T-cell count (* <0.05, ** <0.01, *** <0.001).

To determine whether ongoing antigen recognition by HIV-specific T-cells could be maintaining IFN-γ-producing HIV-specific T-cell responses, we next examined associations between the slopes of change of T-cell response magnitudes between 24 and 168 weeks post-study entry (based on absolute changes on a linear scale) with on-ART virologic parameters, including total cell-associated HIV DNA (CA-DNA), cell-associated HIV RNA (CA-RNA), and plasma HIV RNA by integrase single copy assay (iSCA). The dynamics of responses to the most immunogenic antigens, Gag and Nef (33), along with summed HIV responses, were significantly associated with pre-ART viral loads (**Fig. 4B and Table S5**), despite participants having been on ART for over a median of 7 years when responses were first measured. Strikingly, however, the slopes of change in Nef-specific responses were unique in exhibiting highly significant direct associations with any on-ART virologic parameter after controlling for potential confounding by pre-ART plasma viral load and pre-ART CD4^+^ T-cell count, specifically on-ART CA-DNA (r = 0.496, p = 0.003) and CA-RNA (r = 0.405, p = 0.019) at study entry (**Fig. 4B and Table S5**). These results indicate that both higher frequencies of persistent infected cells (CA-DNA) and higher levels of viral transcription (CA-RNA) were associated with greater maintenance of Nef-specific responses, consistent with ongoing stimulation by infected cells. Slopes of change in HIV-specific T-cell responses were not associated with PD-1 levels on total CD4^+^ or CD8^+^ T-cells (**Fig. 4B and Table S5**), but generally correlated with each other (**Table S6**). Analyzing slopes of change in log_10_-transformed T-cell response magnitudes, reflecting proportional changes in responses rather than absolute changes, revealed significant associations between the dynamics of Nef-specific, Nef/Tat/Rev-specific, and summed HIV-specific T-cell responses with on-ART CA-DNA at study entry (all p<0.05 – **Table S7**), with proportional changes in HIV-specific responses generally correlating with each other (**Table S8**). Thus, whether dynamics were measured on an absolute or proportional change scale, Nef-specific response persistence was uniquely associated with HIV-infected cell frequencies. These results suggest that Nef-specific T-cell responses are preferentially maintained by ongoing interactions with HIV-infected cells, though all responses are likely maintained to some extent by ongoing HIV antigen recognition given their exceptional persistence.

### Recent In Vivo Antigen Recognition by Nef-Specific T-cells

We next investigated whether the functional properties of HIV-specific CD8^+^ T-cells would yield insights into their recent histories of *in vivo* antigen encounter. Data from human studies and animal models have highlighted *ex vivo* granzyme B production as a distinguishing feature of virus-specific effector CD8^+^ T-cells which have recently encountered antigen *in vivo* – either through infection or vaccination (24–28). While granzyme B production can be induced in memory CD8^+^ T-cells, this requires more than 24 hours of *in vitro* stimulation, whereas IFN-γ is produced rapidly from both memory and effector CD8^+^ T-cells (24, 34, 35). Thus, *ex vivo* ELISPOT measurements of granzyme B have been established as an ‘immune diagnostic’ means of identifying effector responses to active infections (34, 36). To quantify the effector functionalities of HIV-specific T-cells on long-term ART, we performed batched granzyme B ELISPOT assays on week 24 and 168 samples (**Fig. 5A**). We focused on the Gag, Pol, and Nef peptide pools, having observed these to be the most immunogenic by IFN-γ ELISPOT. Overall, granzyme B-producing HIV-specific responses were substantially lower in magnitude than IFN-γ responses (**Fig. 5B, 5C and Table S2**). The median magnitudes of granzyme B responses relative to each other were: Nef>Pol>Gag (at both timepoints – **Fig. 5B and Table S2**), contrasting with IFN-γ: Gag>Pol~=Nef (**Fig. 3B and Table S2**). As with IFN-γ, categorizing participants’ granzyme B responses as either increasing, decreasing, or not changing revealed heterogeneity (**Fig. S2**), though proportionally there were fewer decreasing responses, and the population-average levels of granzyme B responses were highly stable over time to all HIV-gene products (**Fig. 5B and Table S4**). In contrast to IFN-γ, we did not observe any significant correlations between the slopes of change of granzyme B responses with virologic measures of HIV persistence (**Tables S9 & S11**). These results may reflect the additional complexity that whereas both IFN-γ–and granzyme B-producing cells can be maintained by infected cells producing antigen, the latter are more likely to also perturb the virologic measures by eliminating infected cells (37).

**Fig. 5.**
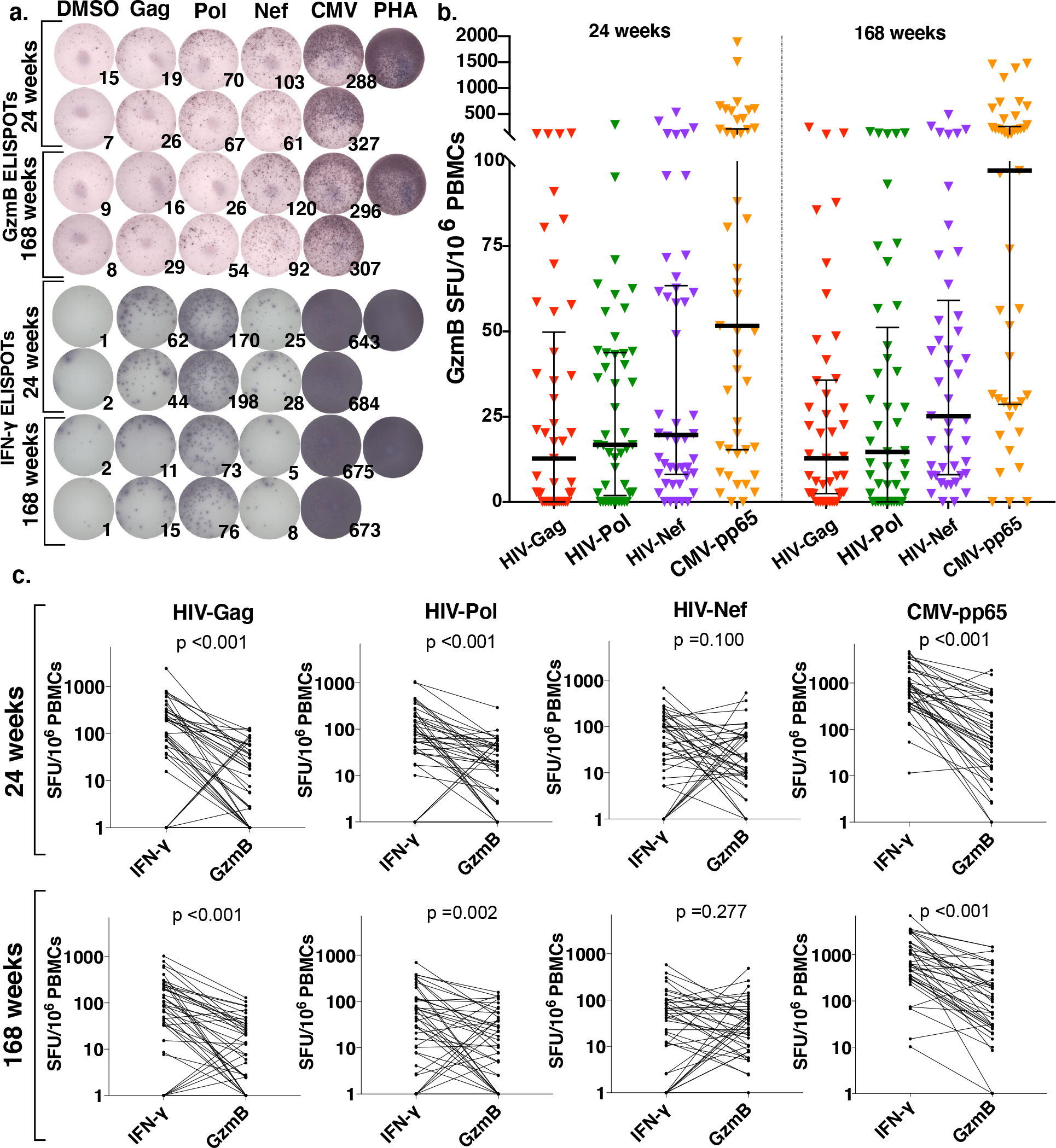
HIV-specific T-cells demonstrate cytotoxic ability, preferentially directed towards HIV-Nef, evidencing recent *in vivo* antigen exposure. **A.** Corresponding granzyme B and IFN-γ ELISPOT results for one participant at both timepoints, with 2×10^5^ PBMCs/well. **B.** Magnitudes of granzyme B responses are shown for two batched on-ART timepoints. Each data point represents the mean number of SFU/10^6^ PBMCs following background subtraction of mean of negative control wells. Vertical lines and error bars represent median and interquartile range for each peptide pool. **C.** Pairwise comparisons of granzyme B versus IFN-γ responses for Gag, Pol, Nef, and CMVpp65 at both timepoints. P-values calculated by Wilcoxon matched pairs signed-rank test.

To further assess the functional profiles of HIV-specific T-cell responses, we performed pairwise comparisons of granzyme B versus IFN-γ responses for each of the gene products tested (**Fig. 5C**). At both timepoints, granzyme B response magnitudes to Gag, Pol, and CMV-pp65 were substantially lower than IFN-γ responses (all p<0.05) (**Fig. 5C**). Contrasting this, the magnitudes of granzyme B versus IFN-γ responses to Nef were not significantly different from each other at either timepoint (p=0.100 at week 24, p=0.277 at week 168). These data indicate that in addition to being preferentially maintained over time, T-cell responses directed against the early HIV gene product Nef disproportionately exhibit effector functional profiles, as compared to the late gene products Gag and Pol (though appreciable granzyme B responses to these gene products were still detected). Persistent HIV-specific granzyme B responses are indicative of recent antigen encounter, supporting the hypothesis that there is *in vivo* stimulation by HIV-infected cells despite suppressive ART.

## Discussion

An important aspect of how HIV persists in individuals on long-term ART is through the evasion of immune recognition, predominately thought to be achieved through the maintenance of strict viral latency, with an additional aspect of anatomical sequestration. This perception that the reservoir is entirely latent has begun to shift lately, in response both to a new understanding of the dynamic nature of the HIV reservoir (driven by the clonal expansion of infected cells), and to new insights into ongoing viral transcriptional activity on ART (15, 38). To date, however, this has yet to prompt widespread re-consideration of the relationship between the HIV-specific T-cell response and the HIV reservoir. The current study provides evidence which challenges the prevailing model of a lack of reservoir immune surveillance, by indicating a level of ongoing antigenic stimulation of HIV-specific T-cells in ART-suppressed individuals. Nef-specific T-cells stood apart from those of other HIV gene products in this regard, supporting that early gene products (Nef, Tat, and Rev – of which only Nef was appreciably immunogenic [as also seen in other studies (21, 33)]) have lower thresholds to expression in a reactivation setting as compared to late gene products (Gag, Pol, and Env), which are expressed only after a cell has built up sufficient levels of Rev to drive nuclear export of unspliced and singly-spliced viral transcripts (39, 40). The preferential maintenance of Nef-specific T-cells was presented as a hypothesis of the current study based, in part, on our previous observation that Nef-specific T-cells recognized cells reactivated from an *in vitro* latency model prior to recognition by Gag-specific T-cells, or detectable Gag expression (32).

Do our results allow for any inferences into how frequently infected cells are recognized by HIV-specific T-cells *in vivo*? While numerous aspects of complexity introduce caveats to such an analysis (e.g. tissue distributions), our data do allow for side-by-side comparisons between the peripheral blood frequencies of infected cells with antigen-expression potential, and those of HIV-specific T-cells – which may be informative. The median frequency of Nef-specific T-cells at week 24 of our study was 78.5/10^6^ PBMCs, whereas the median total frequency of HIV-infected cells (CA-DNA) was 515.7/10^6^ CD4^+^ T-cells (at week 0), or roughly 103/10^6^ PBMCs. These infected cells, however, predominately contain defective proviruses (41), many of which are likely incapable of expressing antigens (42). It can therefore be reasonably estimated that, in most individuals, Nef-specific T-cells are at least as frequent as infected cells with the potential to express antigen. Our data indicating that the former are influenced by the latter therefore suggest that antigen expression is more likely to be a common versus a rare event *in vivo*, amongst infected cells with this potential. Further study is needed, however, and characterizing the clonal dynamics of HIV-specific T-cells may yield additional insights.

Although latency almost certainly contributes to viral persistence, our findings indicating that HIV reservoirs are not fully hidden from circulating cytotoxic T-cells raise the question of what additional mechanisms may be at play. We first consider the role of immune escape – the process by which HIV evades recognition by acquiring mutations in T-cell epitopes. Immune escape plays a critical role in limiting the overall efficacy of the HIV-specific T-cell response in untreated infection, and HIV reservoirs show clear evidence of past selection, in the form of extensive sequence variation in known T-cell epitopes (29). However, the question at hand pertains to HIV-specific T-cell responses that show evidence of being maintained by recent antigen recognition, indicating that they target epitopes which are intact in at least a portion of the reservoir. Further supporting this idea are the previous observations that: i) the fixation of escape mutations leads to the contraction of corresponding T-cell responses (43), and ii) the substantial majority of HIV-specific T-cells that remain detectable after years of ART target epitopes for which escape is not fixed in corresponding reservoir viruses (44, 45). As with latency, our data do not lead us to contest the idea that the fixation of escape mutations in the reservoir diminishes the overall potential for immune recognition, nor the value of therapeutic strategies to address either of these limitations. However, we are still left with the question of how to reconcile our findings indicating an appreciable level of ongoing *in vivo* recognition of infected cells by cytotoxic (granzyme B) T-cells, with the overall stability of HIV reservoir sizes.

We therefore draw from two recent findings in the field to propose how an HIV reservoir may persist without being fully hidden from circulating cytotoxic T-cells. The first derives from the recent demonstrations that the HIV reservoir is predominately composed of infected T-cells that have undergone clonal expansion (46–48), with different clones dynamically ‘waxing and waning’ over time (48). Thus, HIV-specific T-cells may frequently eliminate infected cells, only to have these replaced by clonal expansion of other reservoir-harboring cells. There have been somewhat conflicting recent reports regarding this possibility – from groups that approached the question from different angles – highlighting the need for further study (42, 49, 50).

Second, we have recently reported that reservoir-harboring cells exhibit intrinsic resistance to T-cell mediated elimination (51), mediated in part by BCL-2 over-expression, which antagonizes perforin/granzyme killing (52). In fact, while it has been generally assumed in our field that the encounter between an antigen-expressing HIV-infected cell and a functional (ex. perforin/granzyme releasing) CD8^+^ T-cell will result in elimination, this overlooks the role of the target cell as an active partner in the killing process. Multiple regulatory mechanisms exist, both in physiological and pathological states, by which target cells determine whether or not to undergo apoptosis, despite receiving a perforin/granzyme hit (53, 54). Thus, one way to resolve our findings with others in the field is to propose that the recognition of HIV-infected cells by HIV-specific cytotoxic T-cells may occur with some frequency *in vivo*, but that this often does not result in target cell elimination. An intriguing possibility is that the combined effects of selection, based on intrinsic susceptibility to CD8^+^ T-cells, and clonal expansion of surviving cells may enable the evolution of a resistant reservoir, paralleling the phenomenon of ‘immunoediting’ in cancer (11). While latency reversal will likely be a critical component of curing HIV infection, our findings raise the hypothesis that – in lieu of an ideal latency reversing agent – reductions in HIV reservoirs may be achievable by boosting immune targeting of existing expression of early gene products (such as Nef, and in a manner that targets non-escaped epitopes,) while enhancing cytotoxic function, limiting clonal expansion, and addressing resistance to cytotoxic T-cells in reservoir-harboring cells.

## Methods

### Study Design

For these observational studies, we evaluated participants from two separate populations. The data in **Fig. 1** were collected on participants diagnosed with HIV infection recruited via convenience sampling through Maple Leaf Medical Clinic in Toronto, Canada. Outliers were not defined or excluded. Participants in this Toronto cohort were virally suppressed for a minimum of 2 years prior to a leukapheresis procedure to collect PBMCs, with no reported ART interruptions or detectable viral loads by a commercial clinical assay. The objective of this first study was to evaluate the total ability of participant’s CD8^+^ T-cells to recognize autologous HIV reservoir viruses, and to compare these results with T-cell responses as measured by IFN-γ ELISPOT. All other data for this manuscript were collected on a longitudinal cohort of participants who initiated ART during chronic HIV infection in AIDS Clinical Trials Group (ACTG) trials for treatment-naïve individuals, and enrolled in the ACTG HIV Reservoirs Cohort Study (A5321) (31). A5321 cohort participants were recruited from 17 clinical research sites in the United States through the ACTG network. IFN-γ ELISPOTs were previously performed using samples from 96 participants at A5321 study entry (32), and a subset of 49 participants were selected from the original 96 for this longitudinal sub-study based on sample availability. All gene products and negative controls were tested in duplicate, with one replicate of PHA positive control. Assays performed under these same conditions have been previously validated in other participant cohorts. Outliers were not defined or excluded. Participants in the current sub-study had follow-up at least every 6 months following study entry, with documented sustained viral suppression (plasma HIV RNA levels <50 copies/mL by commercial assays starting at week 48 on ART and at all subsequent timepoints – **Fig. 2**). One participant had a large viral blip (>1,000 copies/mL) 43 weeks prior to their 168 week A5321 study timepoint, and data was right-censored for this participant after the 24 week A5321 study timepoint. Clinical data and paired plasma and PBMC samples were available from pre-ART and on ART study visits. We measured HIV levels (CA-DNA, CA-RNA, and plasma iSCA) and PD-1 levels (on CD4^+^ and CD8^+^ cells) on samples obtained at A5321 study entry (median 7 years on ART), and plasma HIV RNA levels and CD4^+^ T-cell counts were obtained from pre-ART clinical data. One participant later revoked consent for further testing and was excluded from analysis. We hypothesized *a priori* that the long-term dynamics of T-cell responses to the early HIV gene product Nef (measured by IFN-γ ELISPOT) would be associated with infected cell frequencies.

### Virologic Assays

HIV CA-DNA and CA-RNA were measured by quantitative PCR (qPCR) assays in PBMCs using previously described methods (55). CA-DNA and CA-RNA values per million CD4^+^ T-cells were calculated by dividing the total CA-DNA or CA-RNA copies/million PBMCs (normalized for CCR5 copies measured by qPCR as published (55)) by the CD4^+^ T-cell percentage (x 0.01) reported from the same specimen date or from a CD4^+^ T-cell percentage imputed using linear interpolation from specimen dates before and after the CA-DNA or CA-RNA results. Cell-free HIV RNA was quantified by iSCA in blood plasma (5 mL) (56).

### Immunologic Assays

PBMCs obtained at A5321 study entry were stained with the following monoclonal antibodies to evaluate surface PD-1 expression: CD3 APC-H7, CD4 PC5, CD8 V450, PD-1 (clone M1H4) A488 (all from BD Biosciences, San Diego, California, USA), and Live/Dead Aqua (Invitrogen, Grand Island, New York, USA). Cells were fixed in 1% paraformaldehyde, and analyzed using a BD LSR Fortessa (FACSDiva) within 24 hours after staining. Lymphocytes were identified based upon size and granularity. The lymphocyte population was filtered through side scatter area vs. side scatter height histogram to eliminate doublets from the analysis. Single cells were analyzed using Live/Dead Aqua dye exclusion and then CD4^+^ and CD8^+^ populations were defined based on dual expression with CD3. These two populations were plotted against PD-1. Fluorescence minus one (FMO) controls were used to define the PD-1^+^ T-cell populations.

### Quantitative Viral Outgrowth Assay (QVOA)

Quantitative Viral Outgrowth Assays (QVOA) were performed as previously described (57). Briefly, CD4^+^ T-cells were isolated from PBMCs by negative selection (Easysep, Stemcell Technologies) and plated in serial dilution at either 4 or 6 concentrations (12 wells/concentration, 24-well plates). CD4^+^ T-cells were stimulated with phytohemagglutinin (PHA, 2μg/mL) and irradiated allogeneic HIV-negative PBMCs were added to further induce viral reactivation. MOLT-4/CCR5 cells were added at 24 hours post-stimulation as targets for viral infection. Culture media [RPMI 1640 + 10% FBS + 1% Pen/Strep +50U/mL IL-2 + 10ng/mL IL-15 (R10-50-15)] was changed every 3 days and p24 enzyme-linked immunosorbent assay (ELISA, NCI Frederick) was run on day 14 to identify virus-positive wells. Infectious Units per Million CD4^+^ T-cells (IUPM) values were determined using the Extreme Limiting Dilution Analysis (ELDA) software (http://bioinf.wehi.edu.au/software/elda/) (58). Culture supernatants from virus-positive wells were frozen (−80°C) for future use.

### CD8^+^ T-cell Biosensor Assay

<u>Activation of CD4^+^ T-cell Targets</u>: CD4^+^ T-cells were enriched by negative selection (Easysep, Stemcell Technologies), typically starting from 200×10^6^ PBMCs per study participant. These cells were stimulated with 10μg/ml of anti-CD3 (OKT-3) and anti-CD28 (28.2) antibodies (Ultra-LEAF™, Biolegend) in R10-50-15 for 48 hours. <u>Infections</u>: Cells were then washed, and split into 3 equal aliquots (~4×10^6^ cells each) for infection with either: i) JRCSF ii) autologous virus or iii) mock (nothing). The autologous virus stock was generated by pooling equal volumes of all p24^+^ QVOA wells from that study participant, while the JRCSF stock was generated by transfection of 293T cells with plasmid. All viruses were titrated on TZM-bl cells, and used at a MOI of 0.4. After a 2 hour infection period at 37°C 5% CO_2_, cells were washed and then cultured for 48 hours in R10-50. Levels of infection were monitored every 24 hours by surface staining small aliquots with anti-CD3 Brilliant Violet 785™, CD4 Pacific Blue™ (Biolegend), then permeabilizing (BD Cytofix/Cytoperm™) and staining intracellularly with anti-HIV-Gag Kc57-RD1 (Beckman Coulter), and analyzed on a BD LSRFortessa™ flow cytometer. Infections were harvested for co-culture with CD8^+^ T-cells, when they reached 2-4% Gag^+^ within the CD3^+^ gate.

<u>Co-culture with CD8^+^ T-cells</u>: CD8^+^ T-cells autologous to these CD4^+^ targets were enriched on the day of co-culture from freshly a thawed aliquot of 100×10^6^ PBMCs by negative selection (Easysep™, Stemcell Technologies). Infected and mock-infected CD4^+^ T-cells were washed 5x and then co-cultured with CD8^+^ T-cells at ratios of 5 CD8^+^ T-cells to 1 CD4^+^ T-cell, at a total concentration of 5×10^6^ cells/mL in RPMI 1640 + 10% FBS + 1% Pen/Strep +50U/mL IL-2 (R10-50) with 1/100 anti-CD107a-PE antibody (Biolegend) and 1/1,000 Monensin GolgiStop™ (BD). Cells were incubated for a total of 6 hours, with mixing by pipetting every 30 minutes (to facilitate contacts between antigen-specific CD8^+^ cells and targets). Staining and Analysis: Cells were surface stained with anti-CD3 Brilliant Violet 785™, CD4 Pacific Blue™, CD8 Alexa Fluor^®^ 700, and LIVE/DEAD™ Fixable Aqua dye (Thermofisher). Cells were then washed, permeabilized (BD Cytofix/Cytoperm™), stained intracellularly with anti-HIV-Gag Kc57-RD1 (Beckman Coulter), and analyzed on a BD LSRFortessa™ flow cytometer.

### Peptide Pools

The following sets of consensus HIV clade B 15 amino acid peptides (overlapping by 11 amino acids) were supplied by the NIH AIDS Research and Reference Reagent Program: Gag (cat # 8117), Env (cat # 9480), Pol (cat # 6208), Tat (cat # 5138), Rev (cat # 6445), and Nef (cat # 5189). All peptides were dissolved at 5mg/mL in 12.5% DMSO (Corning), and 87.5% PBS (Gibco). Peptides were pooled into whole gene product peptide pools and adjusted to a final concentration of 20μg/mL/peptide in PBS. A CMV-pp65 PepMix peptide pool (JPT Peptide Technologies) was dissolved separately in DMSO and adjusted to a final concentration of 20μg/mL/peptide in PBS.

### IFN-γ and Granzyme B ELISPOT Assays

Multi-screen IP 96-well PVDF plates (Millipore) were either directly coated with 100μL/well of PBS + 0.5μg/mL primary anti-human IFN-γ antibody (clone 1-D1K, Mabtech) overnight at 4°C, or first primed with 20μL of 35% EtOH/well, and immediately washed 6x with 200μL ddH_2_O and then coated with 100μL/well of PBS + 15μg/mL primary anti-human granzyme B antibody (clone GB10, Mabtech) overnight at 4°C. Granzyme B plates were washed 6x with 200μL PBS and blocked with RPMI 10% FBS (Gibco) (‘R-10’) at 37°C 5% CO_2_. PBMCs were thawed and resuspended in R10 and added to plates at 100,000-200,000 cells/well. HIV peptide pools (20μg/mL/peptide) were added at 10μL/well for a final concentration of 1μg/mL/peptide in <0.5% DMSO. CMV-pp65 peptide pools were added at 10μL/well for a final concentration of 1μg/mL/peptide in <0.5% DMSO. PHA was dissolved in DMSO and PBS to 200μg/mL, and then added to a final concentration of 1μg/mL as a positive control. 0.5% DMSO in PBS and R-10 media were used as negative controls. Plates were incubated for 18 hours at 37°C with 5% CO_2_. Plates were washed 6x with 200μL PBS. Biotinylated secondary IFN-γ antibody (clone 7-B6-1, Mabtech) at 0.5μg/mL in PBS, or biotinylated secondary anti-granzyme B antibody (clone GB11, Mabtech) at 1.0μg/mL in PBS was added to the plates to a final volume of 100μL and incubated for 1 hour in the dark. Plates were then washed 6x with PBS and 0.5μg/mL of Streptavidin-ALP (Mabtech) was added to IFN-γ plates at 100μL/well, and 1μg/mL of Streptavidin-ALP (Mabtech) was added to granzyme B plates at 100μL/well and incubated for 1 hour. Plates were washed 6x with PBS and then color development substrate solution: 10.6mL of ddH_2_O, 400μL 25x AP Color Development Buffer (Biorad), 100μL AP color reagent A (Biorad), and 100μL AP color reagent B (Biorad) was added to the plate at 100μL/well for 15 minutes. After removal of the color development substrate solution, 0.5% of Tween-20 in PBS was added at 100μL/well for 10 minutes. Plates were then washed with water, and left overnight to dry. Plates were counted using Immunospot S6 Ultimate Analyzer and ImmunoSpot software (Cellular Technology Limited).

### Statistics

Statistical analyses including univariate statistics and Spearman *r* correlations and partial correlations (adjusting for potential confounders) were conducted in SAS University Edition. Slopes of change in **Fig. 4B** and **Tables S5-S6 and S9-S10** were calculated based on absolute changes on a linear scale between weeks 24-168 post-A5321 study entry, excluding participants who had a change from 0 magnitude to 0 magnitude. Analyses for **Tables S7-S8 and S11-S12** used slopes of change calculated based on proportional changes on a log_10_ scale between weeks 24-168 post-A5321 study entry, excluding participants who had a change from 0 magnitude to 0 magnitude; slopes reflecting a change from 0 magnitude to a non-zero magnitude were analyzed as the highest rank, and slopes reflecting a change from a non-zero magnitude to 0 magnitude were analyzed as the lowest rank. Statistical analyses including one-way ANOVA and Wilcoxon signed-rank tests were conducted in GraphPad Prism v.8.0. Plots for figures were made in GraphPad Prism v.8.0 and SAS University Edition. A custom code was generated in MATLAB v.9.7 to produce the correlogram in **Fig. 4B**. All linear mixed-effects models were conducted using the R ‘lme4’ package (59), with random intercepts only or both random intercepts and random slopes on the participant level modeled for the random effects, as assessed by significantly improved model fit when random slopes were included (using the R ANOVA function to compare models); multiple comparisons were made where indicated using the R ‘multcomp’ package (60), adjusting for multiple comparisons using Tukey’s all-pair method. Linear mixed-effects models used log_10_-transformed response data, treating zero-valued responses as missing data. Imputation was not used to address missing data, as the degree of missingness was low. All statistical tests were two-sided, α=0.05.

### Study Approval

Ethics oversight for part 1 of this study, for participants from Maple Leaf Medical Clinic, was provided by The George Washington University under IRB protocol #021750. Participants were recruited via convenience sampling by research staff at Maple Leaf Medical Clinic during routine care visits; prospective participants were provided a verbal description of the research and a copy of the informed consent form, which detailed the study’s objectives, risks, and benefits. For part 2 of this study, each ACTG A5321 clinical research site had the A5321 protocol and consent form, and its relevant parental protocols and consent forms, approved by their local IRB, as well as registered with and approved by the Division of AIDS (DAIDS) Regulatory Support Center (RSC) Protocol Registration Office, prior to any participant recruitment and enrollment. Once a participant for study entry was identified, details were carefully discussed with the prospective participant by clinical staff at the site. The participant (or, when necessary, the parent or legal guardian if the participant was under guardianship) was asked to read and sign the ACTG-approved protocol consent form.

## Author contributions

RBJ designed the study. EMS, ARW, RT, AST, SHH, TRD, ST, JKB, TMM, AD, GQL, AG, PK, WDCA, JCC, and BM performed experiments. EMS, ARW, RT, ST, SHH, JKB, CML, RJB, BM, JCC, and JWM analyzed data. RTG, DKM, JJE, and JWM provided participant data. EMS, ARW, and RBJ wrote the manuscript. All authors contributed to the critical revision of the manuscript.

## Acknowledgments

This work was supported by 1) the Martin Delaney ‘BELIEVE’ Collaboratory (NIH grant 1UM1AI26617), which is supported by the following NIH Co-Funding and Participating Institutes and Centers: NIAID, NCI, NICHD, NHLBI, NIDA, NIMH, NIA, FIC, and OAR; 2) by the National Institute of Allergy and Infectious Diseases of the National Institutes of Health under Award Number UM1 AI068634, UM1 AI068636 and UM1 AI106701; and 3) by a grant from the AIDS Clinical Trials Group Network (ACTG) to the University of Pittsburgh Virology Specialty Laboratory. It was also supported in part by the NIH funded R01 grants AI31798 and AI147845, and by an ACTG special projects grant (to RBJ). We thank Shy Genel for assistance with MATLAB coding. We gratefully acknowledge the contributions of the study participants, without whom this work would not be possible. The content is solely the responsibility of the authors and does not necessarily represent the official views of the National Institutes of Health.

JWM is a consultant to Gilead Sciences and Merck, and owns share options in Co-Crystal Pharmaceuticals and Abound Bio, Inc., which are not involved in the current work. JJE has research funding outside the current work from ViiV Healthcare, Gilead Sciences and Janssen, and has consulting income from ViiV Healthcare, Gilead Sciences, Janssen, and Merck. BM has received research funding from Gilead Sciences. RTG has served on a scientific advisory board for Merck. EMS volunteers on the Board of Directors of the non-profit community clinic the Berkeley Community Health Project. The authors declare that they have no other perceived conflicts of interest.

