## Supplemental Data for "HIV-specific T-cell responses reflect substantive in vivo interactions with infected cells despite long-term therapy"

### Supplementary materials

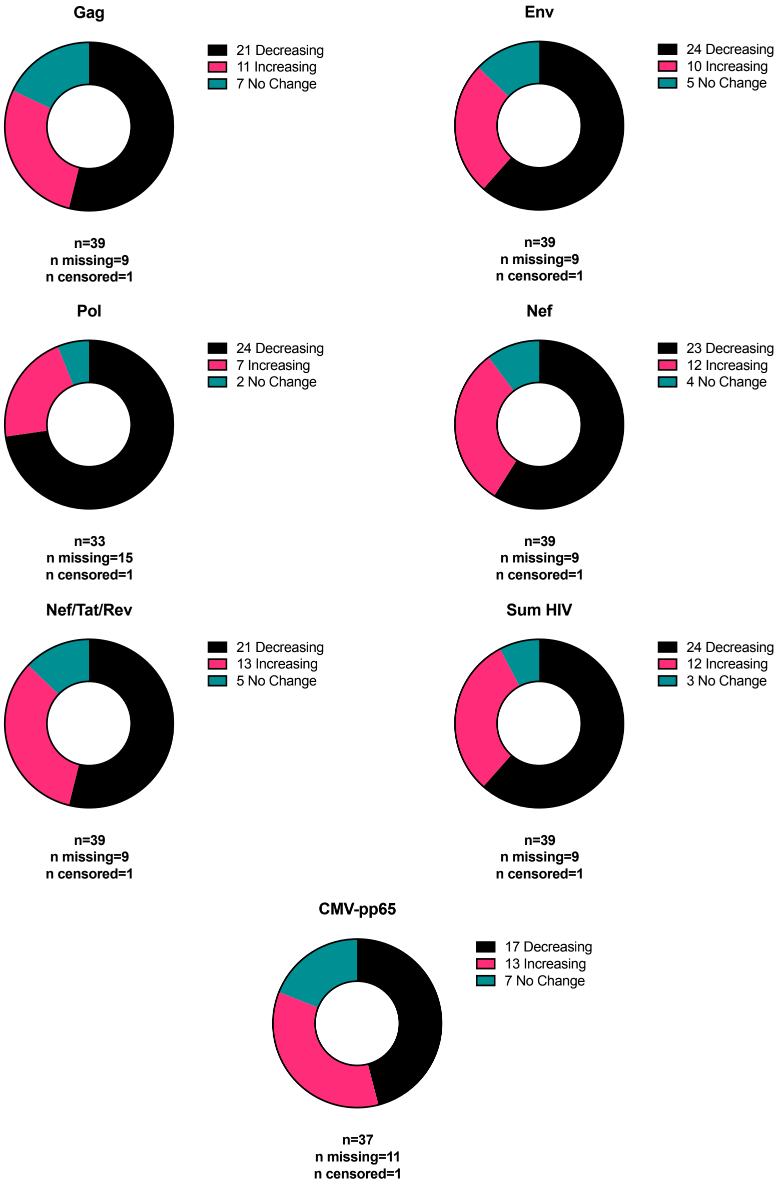


**Figure S1. Patterns of longitudinal IFN-γ T-cell response changes between 24-168 weeks post-A5321 study entry.** Participant T-cell responses were categorized as either increasing, decreasing, or not changing (defined as ≤15% change in either direction) between the two batched on-ART timepoints, and data is shown in parts-of-whole plots. Participants were categorized as missing if they had a missing value for a response at either timepoint. Participants were categorized as censored if they had a censored value for a response at either timepoint.

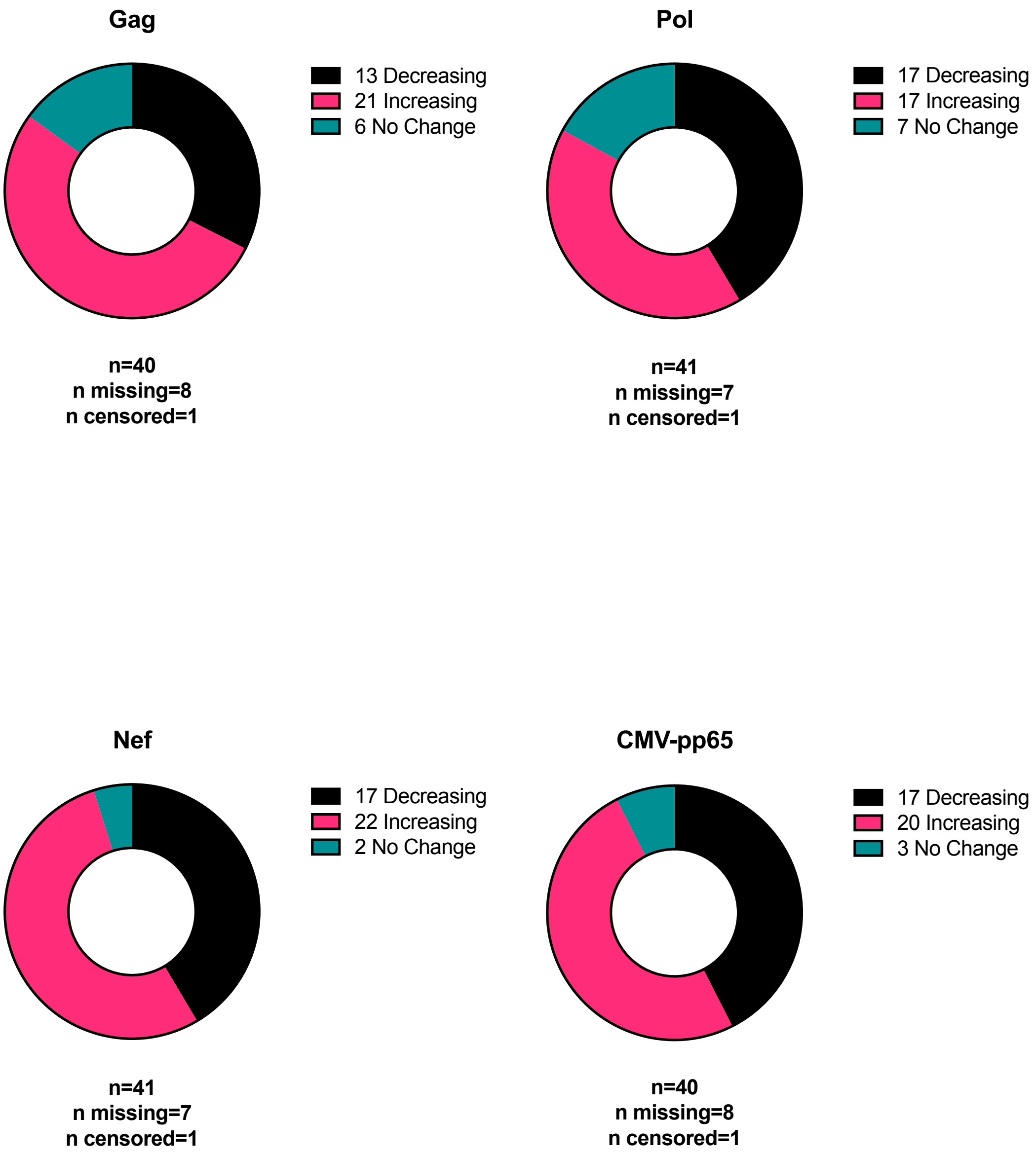


**Figure S2. Patterns of longitudinal granzyme B T-cell response changes between 24-168 weeks post-A5321 study entry.** Participant T-cell responses were categorized as either increasing, decreasing, or not changing (defined as ≤15% change in either direction) between the two batched on-ART timepoints, and data is shown in parts-of-whole plots. Participants were categorized as missing if they had a missing value for a response at either timepoint. Participants were categorized as censored if they had a censored value for a response at either timepoint.

| **Table S1. Demographic and ART-related characteristics of participants for biosensor assay** | | | | | | | | |
|---|---|---|---|---|---|---|---|---|
| *Panel B and C participants (n=11)* | | | | | | | | |
| ***Continuous Variables*** | **Median** | **Range** | | **Missing** | | ***Categorical Variables*** | **n** | **%** |
| | | Lower | Upper | n | % | | | |
| Age at leukapheresis (years) | 49 | 34 | 58 | 1 | 9.09% | ART Initiation Time[A] | | |
| | | | | | | Early/Acute | 2 | 18.18% |
| | | | | | | Late/Chronic | 9 | 81.82% |
| | | | | | | Missing | 0 | 0.00% |
| | | | | | | Race/Ethnicity | | |
| | | | | | | Black (non-Hispanic) | 1 | 9.09% |
| | | | | | | Hispanic (regardless of race) | 2 | 18.18% |
| | | | | | | White (non-Hispanic) | 7 | 63.64% |
| | | | | | | Missing | 1 | 9.09% |
| | | | | | | Sex | | |
| | | | | | | Female | 0 | 0.00% |
| | | | | | | Male | 11 | 100.00% |
| | | | | | | Missing | 0 | 0.00% |
| *Panel D participants (n=9)* | | | | | | | | |
| ***Continuous Variables*** | **Median** | **Range** | | **Missing** | | ***Categorical Variables*** | **n** | **%** |
| | | Lower | Upper | n | % | | | |
| Age at leukapheresis (years) | 50 | 34 | 56 | 1 | 11.11% | ART Initiation Time[A] | | |
| | | | | | | Early/Acute | 1 | 11.11% |
| | | | | | | Late/Chronic | 8 | 88.89% |
| | | | | | | Missing | 0 | 0.00% |
| | | | | | | Race/Ethnicity | | |
| | | | | | | Black (non-Hispanic) | 0 | 0.00% |
| | | | | | | Hispanic (regardless of race) | 1 | 11.11% |
| | | | | | | White (non-Hispanic) | 7 | 77.78% |
| | | | | | | Missing | 1 | 11.11% |
| | | | | | | Sex | | |
| | | | | | | Female | 0 | 0.00% |
| | | | | | | Male | 9 | 100.00% |
| | | | | | | Missing | 0 | 0.00% |

[A]ART initiation early/acute defined as <3 months after known or estimated infection date

**Table S2. Summary statistics for T-cell responses**

| *IFN-γ Responses - Study entry* | | | | | | |
|---|---|---|---|---|---|---|
| **Response** | **Median** | **Q1** | **Q3** | **n** | **n missing** | **n censored** |
| *Gag* | 125.0 | 22.5 | 315.0 | 49 | 0 | 0 |
| *Env* | 20.0 | 0.0 | 45.0 | 49 | 0 | 0 |
| *Pol* | 95.0 | 12.5 | 317.5 | 49 | 0 | 0 |
| *Nef/Tat/Rev* | 67.5 | 17.5 | 265.0 | 49 | 0 | 0 |
| *Sum HIV* | 415.0 | 112.5 | 1087.5 | 49 | 0 | 0 |
| *CMV-pp65* | 1503.8 | 433.8 | 2122.5 | 44 | 5 | 0 |
| *IFN-γ Responses - 24 weeks* | | | | | | |
| **Response** | **Median** | **Q1** | **Q3** | **n** | **n missing** | **n censored** |
| *Gag* | 103.0 | 36.2 | 299.4 | 43 | 6 | 0 |
| *Env* | 22.9 | 0.0 | 105.2 | 43 | 6 | 0 |
| *Pol* | 78.5 | 30.9 | 196.8 | 43 | 6 | 0 |
| *Nef* | 78.5 | 15.4 | 163.6 | 43 | 6 | 0 |
| *Tat* | 5.7 | 0.0 | 38.8 | 43 | 6 | 0 |
| *Rev* | 2.6 | 0.0 | 33.1 | 43 | 6 | 0 |
| *Nef/Tat/Rev* | 69.5 | 17.7 | 146.8 | 43 | 6 | 0 |
| *Sum HIV* | 372.2 | 122.6 | 824.5 | 43 | 6 | 0 |
| *CMV-pp65* | 732.3 | 336.8 | 1750.0 | 43 | 6 | 0 |
| *IFN-γ Responses - 168 weeks* | | | | | | |
| **Response** | **Median** | **Q1** | **Q3** | **n** | **n missing** | **n censored** |
| *Gag* | 87.0 | 32.1 | 233.7 | 44 | 4 | 1 |
| *Env* | 10.3 | 0.0 | 47.7 | 44 | 4 | 1 |
| *Pol* | 44.7 | 6.5 | 112.8 | 44 | 4 | 1 |
| *Nef* | 43.3 | 4.0 | 86.2 | 44 | 4 | 1 |
| *Tat* | 0.0 | 0.0 | 35.5 | 44 | 4 | 1 |
| *Rev* | 0.0 | 0.0 | 15.3 | 44 | 4 | 1 |
| *Nef/Tat/Rev* | 42.2 | 5.4 | 103.2 | 44 | 4 | 1 |
| *Sum HIV* | 250.6 | 117.0 | 517.6 | 44 | 4 | 1 |
| *CMV-pp65* | 610.3 | 301.1 | 1505.3 | 42 | 6 | 1 |

| *GzmB Responses - 24 weeks* | | | | | | |
|---|---|---|---|---|---|---|
| **Response** | **Median** | **Q1** | **Q3** | **n** | **n missing** | **n censored** |
| *Gag* | 12.7 | 0.0 | 43.8 | 45 | 4 | 0 |
| *Pol* | 16.7 | 2.5 | 43.6 | 46 | 3 | 0 |
| *Nef* | 19.6 | 8.3 | 62.6 | 46 | 3 | 0 |
| *CMV-pp65* | 51.6 | 15.4 | 203.3 | 45 | 4 | 0 |
| *GzmB Responses - 168 weeks* | | | | | | |
| **Response** | **Median** | **Q1** | **Q3** | **n** | **n missing** | **n censored** |
| *Gag* | 13.0 | 2.5 | 35.7 | 44 | 4 | 1 |
| *Pol* | 14.8 | 1.3 | 51.1 | 44 | 4 | 1 |
| *Nef* | 27.6 | 9.1 | 59.1 | 44 | 4 | 1 |
| *CMV-pp65* | 102.7 | 28.6 | 263.0 | 44 | 4 | 1 |

**Table S3. Linear mixed-effects model results for differences in HIV-specific T-cell responses between gene products**

| *IFN-γ Responses* [A] | | | | | *GzmB Responses* [A] | | | | |
|---|---|---|---|---|---|---|---|---|---|
| **Comparison** | **Mean Difference** [B] | **95% LCL** | **95% UCL** | **p-Value** [C] | **Comparison** | **Mean Difference** [B] | **95% LCL** | **95% UCL** | **p-Value** [C] |
| *Gag vs. Env* | 0.737 | 0.401 | 1.073 | **<0.001** | *Gag vs. Pol* | -0.095 | -0.330 | 0.139 | 0.602 |
| *Gag vs. Pol* | 0.166 | -0.164 | 0.496 | 0.698 | *Gag vs. Nef* | -0.142 | -0.366 | 0.083 | 0.297 |
| *Gag vs. Nef* | 0.597 | 0.275 | 0.918 | **<0.001** | *Pol vs. Nef* | -0.046 | -0.270 | 0.177 | 0.875 |
| *Gag vs. Tat* | 0.947 | 0.591 | 1.303 | **<0.001** | | | | | |
| *Gag vs. Rev* | 1.083 | 0.726 | 1.441 | **<0.001** | | | | | |
| *Env vs. Pol* | -0.571 | -0.912 | -0.230 | **<0.001** | | | | | |
| *Env vs. Nef* | -0.140 | -0.473 | 0.193 | 0.831 | | | | | |
| *Env vs. Tat* | 0.210 | -0.156 | 0.577 | 0.567 | | | | | |
| *Env vs. Rev* | 0.346 | -0.021 | 0.714 | 0.077 | | | | | |
| *Pol vs. Nef* | 0.431 | 0.104 | 0.758 | **0.003** | | | | | |
| *Pol vs. Tat* | 0.781 | 0.420 | 1.142 | **<0.001** | | | | | |
| *Pol vs. Rev* | 0.917 | 0.556 | 1.279 | **<0.001** | | | | | |
| *Nef vs. Tat* | 0.351 | -0.003 | 0.704 | 0.054 | | | | | |
| *Nef vs. Rev* | 0.487 | 0.132 | 0.841 | **0.002** | | | | | |
| *Tat vs. Rev* | 0.136 | -0.250 | 0.522 | 0.913 | | | | | |

[A]Responses measured at weeks 24 and 168 post-study entry

[B]Least square mean difference in $log_{10}$ ELISPOT spots/$10^6$ PBMCs

[C]Adjusted for multiple comparisons using Tukey's method

LCL - lower confidence limit; UCL - upper confidence limit

| Table S4. Linear mixed-effects model results for overall effect of time on T-cell responses | | | | | |
|---|---|---|---|---|---|
| | | *IFN-γ Responses* | | | |
| **Response[A]** | **Beta[B]** | **95% LCL** | **95% UCL** | **p-Value** | **Mean % Change per Week (ELISPOT spots/$10^6$ PBMCs)** |
| *Gag* | -0.00153 | -0.00277 | -0.00027 | **0.020** | -0.35% |
| *Env* | -0.00271 | -0.00426 | -0.00107 | **0.002** | -0.62% |
| *Pol* | -0.00239 | -0.00367 | -0.00109 | **0.001** | -0.55% |
| *Nef* | -0.00116 | -0.00262 | 0.00031 | 0.125 | n.s. |
| *Nef/Tat/Rev* | -0.00121 | -0.00240 | -0.00002 | 0.052 | n.s. |
| *Sum HIV* | -0.00099 | -0.00212 | 0.00014 | 0.090 | n.s. |
| *CMV-pp65* | -0.00048 | -0.00141 | 0.00046 | 0.319 | n.s. |
| | | *GzmB Responses* | | | |
| **Response[A]** | **Beta[B]** | **95% LCL** | **95% UCL** | **p-Value** | **Mean % Change per Week (ELISPOT spots/$10^6$ PBMCs)** |
| *Gag* | -0.00077 | -0.00249 | 0.00096 | 0.386 | n.s. |
| *Pol* | 0.00011 | -0.00130 | 0.00151 | 0.879 | n.s. |
| *Nef* | 0.00006 | -0.00147 | 0.00157 | 0.938 | n.s. |
| *CMV-pp65* | 0.00145 | 0.00011 | 0.00282 | **0.040** | 0.33% |

[A]Modeling $log_{10}$-transformed magnitudes

[B]Modeling effect of time in weeks

LCL - lower confidence limit; UCL - upper confidence limit

n.s. - not significant

**Table S5. Spearman correlations between slopes of change in magnitudes of IFN-γ T-cell responses from week 24 to week 168 with virologic and immunologic parameters**

| **Variable** | | *Gag* | *Env* | *Pol* | *Nef* | *Nef/Tat/Rev* | *Sum HIV* | *CMV-pp65* |
|---|---|---|---|---|---|---|---|---|
| | r | 0.104 | 0.129 | 0.237 | **0.508** | 0.409 | 0.216 | 0.051 |
| | p-value | 0.546 | 0.461 | 0.158 | **0.001** | 0.015 | 0.186 | 0.762 |
| HIV CA-DNA at A5321 entry (cps/$10^6$ CD4+ T-cells) | n | 36 | 35 | 37 | 37 | 35 | 39 | 37 |
| | Adjusted[A] r | 0.069 | 0.157 | 0.224 | **0.496** | 0.342 | 0.179 | - |
| | Adjusted[A] p-value | 0.698 | 0.383 | 0.195 | **0.003** | 0.051 | 0.290 | - |
| | r | 0.296 | 0.216 | 0.165 | **0.455** | 0.257 | 0.307 | 0.034 |
| | p-value | 0.089 | 0.227 | 0.344 | **0.006** | 0.148 | 0.065 | 0.843 |
| HIV CA-RNA at A5321 entry (cps/$10^6$ CD4+ T-cells) | n | 34 | 33 | 35 | 35 | 33 | 37 | 37 |
| | Adjusted[A] r | 0.211 | 0.199 | 0.154 | **0.405** | 0.222 | 0.252 | - |
| | Adjusted[A] p-value | 0.245 | 0.283 | 0.391 | **0.019** | 0.230 | 0.144 | - |
| | r | -0.096 | -0.302 | -0.019 | 0.158 | -0.013 | -0.121 | 0.129 |
| | p-value | 0.582 | 0.083 | 0.914 | 0.358 | 0.942 | 0.470 | 0.454 |
| HIV plasma RNA via iSCA at A5321 entry (cps/mL) | n | 35 | 34 | 36 | 36 | 34 | 38 | 36 |
| | Adjusted[A] r | -0.325 | -0.424 | -0.098 | -0.104 | -0.153 | -0.346 | - |
| | Adjusted[A] p-value | 0.065 | 0.016 | 0.583 | 0.560 | 0.402 | 0.039 | - |
| | r | 0.034 | 0.292 | 0.018 | 0.008 | 0.050 | 0.059 | -0.226 |
| | p-value | 0.853 | 0.105 | 0.922 | 0.966 | 0.785 | 0.738 | 0.206 |
| %PD-1+ CD4+ cells at A5321 entry | n | 32 | 32 | 33 | 33 | 32 | 35 | 33 |
| | Adjusted[A] r | -0.018 | 0.316 | -0.001 | -0.093 | -0.029 | 0.002 | - |
| | Adjusted[A] p-value | 0.924 | 0.089 | 0.994 | 0.618 | 0.881 | 0.990 | - |
| | r | 0.079 | 0.316 | 0.009 | -0.022 | 0.003 | 0.067 | -0.240 |
| | p-value | 0.667 | 0.078 | 0.960 | 0.902 | 0.989 | 0.704 | 0.179 |
| %PD-1+ CD8+ cells at A5321 entry | n | 32 | 32 | 33 | 33 | 32 | 35 | 33 |
| | Adjusted[A] r | 0.018 | 0.330 | -0.015 | -0.137 | -0.070 | 0.007 | - |
| | Adjusted[A] p-value | 0.927 | 0.075 | 0.937 | 0.463 | 0.713 | 0.971 | - |
| | r | **0.435** | 0.230 | 0.133 | **0.472** | 0.216 | **0.374** | 0.251 |
| | p-value | **0.008** | 0.183 | 0.431 | **0.003** | 0.213 | **0.019** | 0.134 |
| Pre-ART plasma HIV-1 RNA ($log_{10}$cps/mL) | n | 36 | 35 | 37 | 37 | 35 | 39 | 37 |
| | Adjusted[B] r | **0.428** | 0.216 | 0.112 | **0.479** | 0.166 | **0.361** | - |
| | Adjusted[B] p-value | **0.010** | 0.219 | 0.516 | **0.003** | 0.348 | **0.026** | - |
| | r | -0.105 | 0.081 | -0.080 | -0.291 | -0.259 | -0.176 | **-0.426** |
| Pre-ART CD4+ T-cell count (cells/$mm^3$) | p-value | 0.544 | 0.644 | 0.638 | 0.080 | 0.132 | 0.283 | **0.009** |
| | n | 36 | 35 | 37 | 37 | 35 | 39 | 37 |

[A]Controlling for Pre-ART plasma HIV-1 RNA ($log_{10}$cps/mL) and Pre-ART CD4+ T-cell count (cells/$mm^3$)

[B]Controlling for HIV CA-DNA at A5321 entry (cps/$10^6$ CD4+ T-cells)

Note: zero-value slopes reflecting a change from 0 magnitude to 0 magnitude excluded

**Table S6. Spearman correlations between slopes of change in magnitudes of IFN-γ T-cell responses from week 24 to week 168**

| | | *Gag* | *Env* | *Pol* | *Nef* | *Nef/Tat/Rev* | *Sum HIV* | *CMV-pp65* |
|---|---|---|---|---|---|---|---|---|
| *Gag* | r | - | | | | | | |
| | p-value | - | | | | | | |
| | n | - | | | | | | |
| *Env* | r | **0.616** | - | | | | | |
| | p-value | **<0.001** | - | | | | | |
| | n | 33 | - | | | | | |
| *Pol* | r | **0.573** | **0.365** | - | | | | |
| | p-value | **<0.001** | **0.037** | - | | | | |
| | n | 35 | 33 | - | | | | |
| *Nef* | r | **0.557** | **0.442** | **0.447** | - | | | |
| | p-value | **0.001** | **0.010** | **0.006** | - | | | |
| | n | 35 | 33 | 36 | - | | | |
| *Nef/Tat/Rev* | r | **0.552** | **0.361** | **0.544** | **0.698** | - | | |
| | p-value | **0.001** | **0.046** | **0.001** | **<0.001** | - | | |
| | n | 33 | 31 | 34 | 34 | - | | |
| *Sum HIV* | r | **0.922** | **0.662** | **0.711** | **0.688** | **0.726** | - | |
| | p-value | **<0.001** | **<0.001** | **<0.001** | **<0.001** | **<0.001** | - | |
| | n | 36 | 35 | 37 | 37 | 35 | - | |
| *CMV-pp65* | r | 0.319 | 0.331 | 0.263 | **0.515** | **0.455** | **0.434** | - |
| | p-value | 0.066 | 0.060 | 0.127 | **0.002** | **0.008** | **0.007** | - |
| | n | 34 | 33 | 35 | 35 | 33 | 37 | - |

Note: zero-value slopes reflecting a change from 0 magnitude to 0 magnitude excluded

**Table S7. Spearman correlations between slopes of change in $log_{10}$-magnitudes of IFN-γ T-cell responses from week 24 to week 168 with virologic and immunologic parameters**

| Variable | | *Gag* | *Env* | *Pol* | *Nef* | *Nef/Tat/Rev* | *Sum HIV* | *CMV-pp65* |
|---|---|---|---|---|---|---|---|---|
| HIV CA-DNA at A5321 entry (cps/$10^6$ CD4+ T-cells) | r | 0.204 | 0.071 | 0.231 | **0.505** | **0.385** | **0.377** | 0.009 |
| | p-value | 0.233 | 0.685 | 0.169 | **0.001** | **0.022** | **0.018** | 0.956 |
| | n | 36 | 35 | 37 | 37 | 35 | 39 | 37 |
| | Adjusted[A] r | 0.222 | 0.147 | 0.260 | **0.481** | **0.355** | **0.358** | - |
| | Adjusted[A] p-value | 0.208 | 0.414 | 0.131 | **0.003** | **0.043** | **0.030** | - |
| HIV CA-RNA at A5321 entry (cps/$10^6$ CD4+ T-cells) | r | 0.186 | 0.049 | -0.084 | 0.333 | 0.057 | 0.345 | 0.020 |
| | p-value | 0.291 | 0.785 | 0.631 | 0.051 | 0.751 | 0.037 | 0.907 |
| | n | 34 | 33 | 35 | 35 | 33 | 37 | 37 |
| | Adjusted[A] r | 0.182 | 0.044 | -0.126 | 0.275 | 0.022 | 0.312 | - |
| | Adjusted[A] p-value | 0.318 | 0.812 | 0.485 | 0.121 | 0.908 | 0.068 | - |
| HIV plasma RNA via iSCA at A5321 entry (cps/mL) | r | -0.177 | -0.172 | 0.178 | 0.229 | 0.038 | -0.120 | 0.120 |
| | p-value | 0.309 | 0.330 | 0.298 | 0.178 | 0.829 | 0.474 | 0.487 |
| | n | 35 | 34 | 36 | 36 | 34 | 38 | 36 |
| | Adjusted[A] r | -0.216 | -0.226 | 0.116 | 0.078 | -0.067 | -0.253 | - |
| | Adjusted[A] p-value | 0.228 | 0.214 | 0.513 | 0.661 | 0.715 | 0.136 | - |
| %PD-1+ CD4+ cells at A5321 entry | r | 0.004 | 0.165 | 0.094 | 0.077 | 0.017 | 0.216 | -0.193 |
| | p-value | 0.984 | 0.367 | 0.604 | 0.670 | 0.926 | 0.213 | 0.282 |
| | n | 32 | 32 | 33 | 33 | 32 | 35 | 33 |
| | Adjusted[A] r | -0.012 | 0.182 | 0.033 | -0.024 | -0.042 | 0.165 | - |
| | Adjusted[A] p-value | 0.950 | 0.336 | 0.860 | 0.897 | 0.824 | 0.360 | - |
| %PD-1+ CD8+ cells at A5321 entry | r | 0.085 | 0.203 | 0.129 | 0.052 | -0.067 | 0.247 | -0.170 |
| | p-value | 0.642 | 0.265 | 0.474 | 0.774 | 0.717 | 0.153 | 0.343 |
| | n | 32 | 32 | 33 | 33 | 32 | 35 | 33 |
| | Adjusted[A] r | 0.067 | 0.211 | 0.064 | -0.055 | -0.125 | 0.198 | - |
| | Adjusted[A] p-value | 0.726 | 0.263 | 0.732 | 0.770 | 0.510 | 0.269 | - |
| Pre-ART plasma HIV-1 RNA ($log_{10}$cps/mL) | r | 0.060 | 0.151 | 0.224 | 0.327 | 0.205 | 0.215 | 0.209 |
| | p-value | 0.728 | 0.387 | 0.183 | 0.049 | 0.237 | 0.189 | 0.215 |
| | n | 36 | 35 | 37 | 37 | 35 | 39 | 37 |
| | Adjusted[B] r | 0.038 | 0.143 | 0.206 | 0.309 | 0.156 | 0.188 | - |
| | Adjusted[B] p-value | 0.830 | 0.420 | 0.228 | 0.067 | 0.377 | 0.260 | - |
| Pre-ART CD4+ T-cell count (cells/$mm^3$) | r | 0.034 | 0.222 | 0.033 | -0.231 | -0.144 | -0.138 | -0.318 |
| | p-value | 0.843 | 0.199 | 0.847 | 0.168 | 0.409 | 0.401 | 0.055 |
| | n | 36 | 35 | 37 | 37 | 35 | 39 | 37 |

[A]Controlling for Pre-ART plasma HIV-1 RNA ($log_{10}$cps/mL) and Pre-ART CD4+ T-cell count (cells/$mm^3$)

[B]Controlling for HIV CA-DNA at A5321 entry (cps/$10^6$ CD4+ T-cells)

Note 1: zero-value slopes reflecting a change from 0 magnitude to 0 magnitude excluded

Note 2: slopes reflecting a change from 0 magnitude to a non-zero magnitude set to highest rank

Note 3: slopes reflecting a change from a non-zero magnitude to 0 magnitude set to lowest rank

**Table S8. Spearman correlations between slopes of change in $log_{10}$-magnitudes of IFN-γ T-cell responses from week 24 to week 168**

| | | *Gag* | *Env* | *Pol* | *Nef* | *Nef/Tat/Rev* | *Sum HIV* | *CMV-pp65* |
|---|---|---|---|---|---|---|---|---|
| *Gag* | r | - | | | | | | |
| | p-value | - | | | | | | |
| | n | - | | | | | | |
| *Env* | r | **0.472** | - | | | | | |
| | p-value | **0.006** | - | | | | | |
| | n | 33 | - | | | | | |
| *Pol* | r | **0.535** | **0.425** | - | | | | |
| | p-value | **0.001** | **0.014** | - | | | | |
| | n | 35 | 33 | - | | | | |
| *Nef* | r | **0.489** | 0.304 | **0.414** | - | | | |
| | p-value | **0.003** | 0.086 | **0.012** | - | | | |
| | n | 35 | 33 | 36 | - | | | |
| *Nef/Tat/Rev* | r | **0.387** | **0.421** | **0.541** | **0.587** | - | | |
| | p-value | **0.026** | **0.018** | **0.001** | **<0.001** | - | | |
| | n | 33 | 31 | 34 | 34 | - | | |
| *Sum HIV* | r | **0.896** | **0.657** | **0.642** | **0.606** | **0.600** | - | |
| | p-value | **<0.001** | **<0.001** | **<0.001** | **<0.001** | **<0.001** | - | |
| | n | 36 | 35 | 37 | 37 | 35 | - | |
| *CMV-pp65* | r | 0.105 | 0.277 | 0.084 | 0.237 | **0.386** | 0.161 | - |
| | p-value | 0.555 | 0.118 | 0.633 | 0.170 | **0.027** | 0.343 | - |
| | n | 34 | 33 | 35 | 35 | 33 | 37 | - |

Note 1: zero-value slopes reflecting a change from 0 magnitude to 0 magnitude excluded
Note 2: slopes reflecting a change from 0 magnitude to a non-zero magnitude set to highest rank
Note 3: slopes reflecting a change from a non-zero magnitude to 0 magnitude set to lowest rank

**Table S9. Spearman correlations between slopes of change in magnitudes of GzmB T-cell responses from week 24 to week 168 with virologic and immunologic parameters**

| Variable | | *Gag* | *Pol* | *Nef* | *CMV-pp65* |
|---|---|---|---|---|---|
| HIV CA-DNA at A5321 entry (cps/$10^6$ CD4+ T-cells) | r | -0.196 | -0.273 | -0.141 | 0.093 |
| | p-value | 0.253 | 0.102 | 0.380 | 0.570 |
| | n | 36 | 37 | 41 | 40 |
| | Adjusted[A] r | -0.176 | -0.255 | -0.107 | - |
| | Adjusted[A] p-value | 0.319 | 0.140 | 0.516 | - |
| HIV CA-RNA at A5321 entry (cps/$10^6$ CD4+ T-cells) | r | 0.173 | 0.066 | -0.076 | **0.414** |
| | p-value | 0.319 | 0.702 | 0.643 | **0.009** |
| | n | 35 | 36 | 40 | 39 |
| | Adjusted[A] r | 0.188 | 0.087 | -0.093 | - |
| | Adjusted[A] p-value | 0.294 | 0.625 | 0.580 | - |
| HIV plasma RNA via iSCA at A5321 entry (cps/mL) | r | -0.320 | -0.083 | 0.022 | -0.049 |
| | p-value | 0.065 | 0.634 | 0.895 | 0.769 |
| | n | 34 | 35 | 39 | 38 |
| | Adjusted[A] r | -0.303 | -0.011 | 0.058 | - |
| | Adjusted[A] p-value | 0.092 | 0.954 | 0.732 | - |
| %PD-1+ CD4+ cells at A5321 entry | r | 0.219 | -0.055 | -0.113 | 0.038 |
| | p-value | 0.228 | 0.765 | 0.512 | 0.831 |
| | n | 32 | 32 | 36 | 35 |
| | Adjusted[A] r | 0.231 | -0.050 | -0.102 | - |
| | Adjusted[A] p-value | 0.219 | 0.792 | 0.565 | - |
| %PD-1+ CD8+ cells at A5321 entry | r | 0.289 | -0.006 | -0.011 | 0.008 |
| | p-value | 0.109 | 0.975 | 0.949 | 0.964 |
| | n | 32 | 32 | 36 | 35 |
| | Adjusted[A] r | 0.300 | -0.006 | -0.006 | - |
| | Adjusted[A] p-value | 0.107 | 0.975 | 0.974 | - |
| Pre-ART plasma HIV-1 RNA ($log_{10}$cps/mL) | r | -0.086 | -0.142 | -0.027 | -0.008 |
| | p-value | 0.618 | 0.401 | 0.866 | 0.960 |
| | n | 36 | 37 | 41 | 40 |
| | Adjusted[B] r | -0.025 | -0.072 | 0.007 | - |
| | Adjusted[B] p-value | 0.885 | 0.675 | 0.967 | - |
| Pre-ART CD4+ T-cell count (cells/$mm^3$) | r | 0.041 | 0.003 | 0.176 | 0.039 |
| | p-value | 0.813 | 0.986 | 0.271 | 0.810 |
| | n | 36 | 37 | 41 | 40 |

[A]Controlling for Pre-ART plasma HIV-1 RNA ($log_{10}$cps/mL) and Pre-ART CD4+ T-cell count (cells/$mm^3$)

[B]Controlling for HIV CA-DNA at A5321 entry (cps/$10^6$ CD4+ T-cells)

Note: zero-value slopes reflecting a change from 0 magnitude to 0 magnitude excluded

**Table S10. Spearman correlations between slopes of change in magnitudes of GzmB T-cell responses from week 24 to week 168**

| | | *Gag* | *Pol* | *Nef* | *CMV-pp65* |
|---|---|---|---|---|---|
| *Gag* | r | - | | | |
| | p-value | - | | | |
| | n | - | | | |
| *Pol* | r | **0.595** | - | | |
| | p-value | **<0.001** | - | | |
| | n | 33 | - | | |
| *Nef* | r | **0.360** | **0.414** | - | |
| | p-value | **0.031** | **0.011** | - | |
| | n | 36 | 37 | - | |
| *CMV-pp65* | r | **0.382** | **0.419** | **0.541** | - |
| | p-value | **0.022** | **0.011** | **<0.001** | - |
| | n | 36 | 36 | 40 | - |

Note: zero-value slopes reflecting a change from 0 magnitude to 0 magnitude excluded

**Table S11. Spearman correlations between slopes of change in $\log_{10}$-magnitudes of GzmB T-cell responses from week 24 to week 168 with virologic and immunologic parameters**

| Variable | | *Gag* | *Pol* | *Nef* | *CMV-pp65* |
|---|---|---|---|---|---|
| HIV CA-DNA at A5321 entry (cps/$10^6$ CD4+ T-cells) | r | 0.124 | -0.225 | -0.124 | 0.016 |
| | p-value | 0.472 | 0.180 | 0.441 | 0.922 |
| | n | 36 | 37 | 41 | 40 |
| | Adjusted[A] r | 0.136 | -0.234 | -0.085 | - |
| | Adjusted[A] p-value | 0.442 | 0.176 | 0.605 | - |
| HIV CA-RNA at A5321 entry (cps/$10^6$ CD4+ T-cells) | r | 0.271 | -0.031 | -0.069 | 0.296 |
| | p-value | 0.115 | 0.857 | 0.673 | 0.067 |
| | n | 35 | 36 | 40 | 39 |
| | Adjusted[A] r | 0.263 | -0.043 | -0.088 | - |
| | Adjusted[A] p-value | 0.140 | 0.811 | 0.601 | - |
| HIV plasma RNA via iSCA at A5321 entry (cps/mL) | r | **-0.405** | -0.087 | 0.022 | -0.118 |
| | p-value | **0.018** | 0.619 | 0.894 | 0.480 |
| | n | 34 | 35 | 39 | 38 |
| | Adjusted[A] r | **-0.459** | -0.052 | 0.057 | - |
| | Adjusted[A] p-value | **0.008** | 0.774 | 0.737 | - |
| %PD-1+ CD4+ cells at A5321 entry | r | 0.251 | -0.057 | -0.091 | 0.064 |
| | p-value | 0.165 | 0.757 | 0.597 | 0.714 |
| | n | 32 | 32 | 36 | 35 |
| | Adjusted[A] r | 0.258 | -0.047 | -0.075 | - |
| | Adjusted[A] p-value | 0.169 | 0.806 | 0.673 | - |
| %PD-1+ CD8+ cells at A5321 entry | r | 0.305 | -0.047 | -0.019 | 0.076 |
| | p-value | 0.090 | 0.798 | 0.913 | 0.666 |
| | n | 32 | 32 | 36 | 35 |
| | Adjusted[A] r | 0.309 | -0.040 | -0.010 | - |
| | Adjusted[A] p-value | 0.097 | 0.834 | 0.954 | - |
| Pre-ART plasma HIV-1 RNA ($\log_{10}$cps/mL) | r | 0.053 | -0.012 | -0.027 | 0.022 |
| | p-value | 0.761 | 0.943 | 0.866 | 0.894 |
| | n | 36 | 37 | 41 | 40 |
| | Adjusted[B] r | 0.014 | 0.054 | 0.002 | - |
| | Adjusted[B] p-value | 0.937 | 0.757 | 0.988 | - |
| Pre-ART CD4+ T-cell count (cells/$mm^3$) | r | 0.077 | <-0.001 | 0.194 | 0.048 |
| | p-value | 0.655 | 0.999 | 0.224 | 0.767 |
| | n | 36 | 37 | 41 | 40 |

[A]Controlling for Pre-ART plasma HIV-1 RNA ($\log_{10}$cps/mL) and Pre-ART CD4+ T-cell count (cells/$mm^3$)

[B]Controlling for HIV CA-DNA at A5321 entry (cps/$10^6$ CD4+ T-cells)

Note 1: zero-value slopes reflecting a change from 0 magnitude to 0 magnitude excluded

Note 2: slopes reflecting a change from 0 magnitude to a non-zero magnitude set to highest rank

Note 3: slopes reflecting a change from a non-zero magnitude to 0 magnitude set to lowest rank

**Table S12. Spearman correlations between slopes of change in $log_{10}$-magnitudes of GzmB T-cell responses from week 24 to week 168**

| | | *Gag* | *Pol* | *Nef* | *CMV-pp65* |
|---|---|---|---|---|---|
| *Gag* | r | - | | | |
| | p-value | - | | | |
| | n | - | | | |
| *Pol* | r | 0.264 | - | | |
| | p-value | 0.137 | - | | |
| | n | 33 | - | | |
| *Nef* | r | **0.426** | **0.546** | - | |
| | p-value | **0.010** | **0.001** | - | |
| | n | 36 | 37 | - | |
| *CMV-pp65* | r | **0.582** | **0.537** | **0.628** | - |
| | p-value | **<0.001** | **<0.001** | **<0.001** | - |
| | n | 36 | 36 | 40 | - |

Note 1: zero-value slopes reflecting a change from 0 magnitude to 0 magnitude excluded
Note 2: slopes reflecting a change from 0 magnitude to a non-zero magnitude set to highest rank
Note 3: slopes reflecting a change from a non-zero magnitude to 0 magnitude set to lowest rank
